# Intraspecific competition for a nest and its implication for the fitness of relocating ant colonies

**DOI:** 10.1101/2023.01.24.525355

**Authors:** Eshika Halder, Sumana Annagiri

**Affiliations:** Department of Biological Sciences, Indian Institute of Science Education and Research Kolkata, Mohanpur, West Bengal, 741246, India

**Author notes:** **Address for correspondence:** Behaviour and Ecology Lab, Department of Biological Sciences, Indian Institute of Science Education and Research Kolkata, Mohanpur, West Bengal 741246, India.

**Keywords:** *Diacamma indicum*, Tandem running recruitment, Gamergate, Brood stealing, Aggression

## Abstract

Competitive interaction is important in structuring species composition of a habitat. Several studies have been conducted on intraspecific competition but little is known in the context of the goal-oriented task of colony relocation. Current study examines how relocating colonies respond when the new nest is a limiting resource. Examining this competition across equal sized colonies (n=17) and unequal sized colonies (n=14), we found that most trails had a clear outcome with one of the colony’s occupying the new nest, while 25% of colonies merged. Larger colonies had a significantly higher chance of occupying the new nest and outcompeted the smaller colonies in most of the cases. Colonies that had lower latency to discover and more explorers had a significantly higher chance of gaining control of the new nest and interesting the level of aggression shown by both the competing colonies was comparable. Spatio-temporal analysis of the aggression revealed that the area surrounding the old, new nest and the time at which transportation occurs has higher levels of aggression. While both the competing colonies stole pupa from each other, the mean number of successful stealing by the larger colonies was 8 times larger than smaller colonies while stealing was comparable between similar sized colonies. Relocating colonies experienced significantly more mortality as compared to controls and competition for the new nest imposed additional mortality on losing colonies over the short time. Due to the potential merging of colonies, incorporation of stolen brood and increased mortality, the overall fitness of the colonies is likely to be negatively impacted due to intraspecific competition for a new nest.

## Introduction

Interaction with the environment and other species is an essential component of an ecosystem. These interactions can be both positive and enhance the organism’s fitness (for example mutualism and commensalism) or negative and reduce the organism’s fitness (for example competition, predation and parasitism) and the balance between these interactions play an important role in maintaining the structure and biodiversity within ecological communities. (1,2). Evidently, intraspecific competition results into territoriality issues, nest overdispersion, reduction in colony size and redistribution of labor in response to a limited resource (3). When two organisms compete against each other for a limited resource such as food (4), shelter (5), or mates, there is typically a winner who succeeds in obtaining the limited resource and the loser who is unable to obtain that particular resource (6–8). These competitions can occur through different behavioural displays like ritualistic displays, aggression or visual and/or chemical signaling (9,10). Different animals use various behavioral mechanisms and execute multiple strategies to gain access and control resources. Exploration is one of the major factors that influences the outcome of these competitions, as faster discovery can help organisms to outcompete other individuals (11,12). Sometimes, involvement in aggressive behaviors shifts the competitors to far away area and reduces the chances of encounters in the future (13). In some cases, the lar ger group outcompetes the smaller group as they can distribute their group members over multiple resources whereas the smaller group might not even be able to take over a single resource (14,15).

In social insects such as ants, conflict occurs between the same species or different species leading to mutual aggression (16). The more similar the species’ niche breadth, the stronger will be the competition (17). Hence, competition and aggressiveness have become a ubiquitous phenomenon in ant colonies and become one of the factors for invasion and ecological change (18). Like other organisms, they show aggressive behaviors towards their non-nestmates, which is one of the important components of their competitive ability (19). The level of aggression depends upon the type and cause of interaction such as food availability (20), closeness between competitors (21), resource value (22), type of habitat (23) and seasonal variation (24). They aggressively defend the immediate territory around their nest against other competing ant colonies (13, 22) and aggressiveness directly depends upon the foraging activity of ants (26). Some ants like *Pristomyrmex punctatus* act more aggressively in their foraging sites as compared to the territory of their own nest (27) but on the other hand, *Temnothorax longispinosus* are more aggressive in the vicinity of their own nest (28). Competition and aggression can be expensive in terms of energy expenditure, time loss, possible injury, and mortality (16).

Nest relocation occurs in social insects. Ant colonies relocate from one nest to another due to various environmental stresses, physical damage, destruction of the nest, to groom immature young ants, to store resources such as food and also for the protection against predators (16). Studies on relocation in ants has been conducted in various parts of the world It is documented in temperate woodland of North America (29), India (30), Japan (31), Europe (32) and the Central American rainforest (33,34). There is more than one colony residing in the same area in the environment (35). The intra-specific competition which influences the relocation of ants are resource depletion (36), microclimatic changes (37), to escape from predators and from the infection of parasites and pathogens (38). The process of relocation involves a lot of disruption of daily activities, additional work and a huge risk as the whole colony becomes more vulnerable during the process, further decisions regarding the optimal new nest has to be taken and colony cohesion has to be maintained through the transport of every adult and brood of the colony. (39,40). It is a goal-oriented task that has a defined initiation and termination point.

When there is a large-scale disturbance in the habitat, like flooding many colonies in the region are bound to get disturbed and are likely to search for new nests and relocate – a scenario that is common during the monsoons.

This study examines the intraspecific competition for nests during the relocation in *Diacamma indicum. D. indicum* is 1 cm long ant that belongs to the family Formicidae and subfamily Ponerine. It is found in India, Bangladesh, Sri Lanka, and possibly Japan. They are monomorphic with average colony sizes being 85.35 (standard deviation is 38.79). Colonies contain a single reproductive individual is called gamergate and she can be distinguished by a pair of thoracic appendages termed gemma (41). These ants are known to relocate using the process called tandem running. In this process, some individuals of the colony have information about the target nest and become tandem leaders and lead their nestmates, known as followers, to the target nest one at a time by maintaining physical contact with each other (30).

In this study, we wanted to instigate competition between two relocating colonies for a new nest and investigate the enfolding aggression and its impact on relocation dynamics. As colony size can play a critical role in this competition, we conducted two sperate sets of experiments with similar sized and unequal sized colonies. The specific objective of this study was to determine a) the outcome of intraspecific competition for the new nest when it is a limited resource, b) the factors that causes a colony to win the resource, and c) the fitness consequences of relocation and competition to the winning and losing colony.

## Materials and methods

### Colony collection and maintenance

All the colonies of *Diacamma indicum* were collected from Nadia, West Bengal 23.4710º N, 88.5565º E, from October 2021 to May 2022 using the nest flooding method (42). Colonies were kept in plastic boxes and were provided with *ad libitum* water and food consisting of ant cakes (43). Each colony of *D. indicum* consists of a reproductive individual called gamergate, who was distinguished by the presence of a pair of gemma at the second thoracic segment, several female workers and brood at different stages of development. Each adult female was given unique identity by marking one or more of their body parts (1^st^, 2^nd^ thoracic segment and abdomen) with non-toxic enamel paint. (Testors, Rockford, IL USA) (44). Two neighboring colonies within 5 m distance were used for a single experiment.

### Experimental setup

All the relocation experiments were performed in the laboratory in a 122cm X 92cm X 18 cm arena filled with water to restrict the movements of ants and the arena walls were coated with petroleum jelly (Vaseline) to further prevent ants from escaping the arena. A maze was created with plastic pipes (Precision, India) and placed in the middle of the arena. Each arm of the maze was 15cm in length. The maze was divided into five areas named A, B, C, D and E as depicted in the setup (Figure 1). Each area consists of three arms for example area A was divided into three arms and termed as A1, A2 and A3. Other areas were also divided in a similar way. Two colonies in their old nests (termed ON) were placed at end of arms A2 and B2, and one empty nest, referred to as a new nest (NN), was placed at the end of arm E3. The total distance between the old and new nest is 60 cm. In their natural habitat, relocating colonies traveled distances in the range of 60 cm to 678 cm (42), thus the distance over which colonies relocated in this setup was realistic. Further, neighboring colonies have been seen inhabiting nests as close as 30cm from each other in the natural habitat, in this setup the traveling distance between the two colonies was 60 cm and hence not unlike what occurs in the natural habitat. Relocation was initiated by removing the roof of old nest from both the colonies and placing a white light on the top of both the old nests simultaneously. The new nest consisted of a nest that was identical to the old nest, except that it had an intact roof and a clear nest entrance.

**Figure 1.**
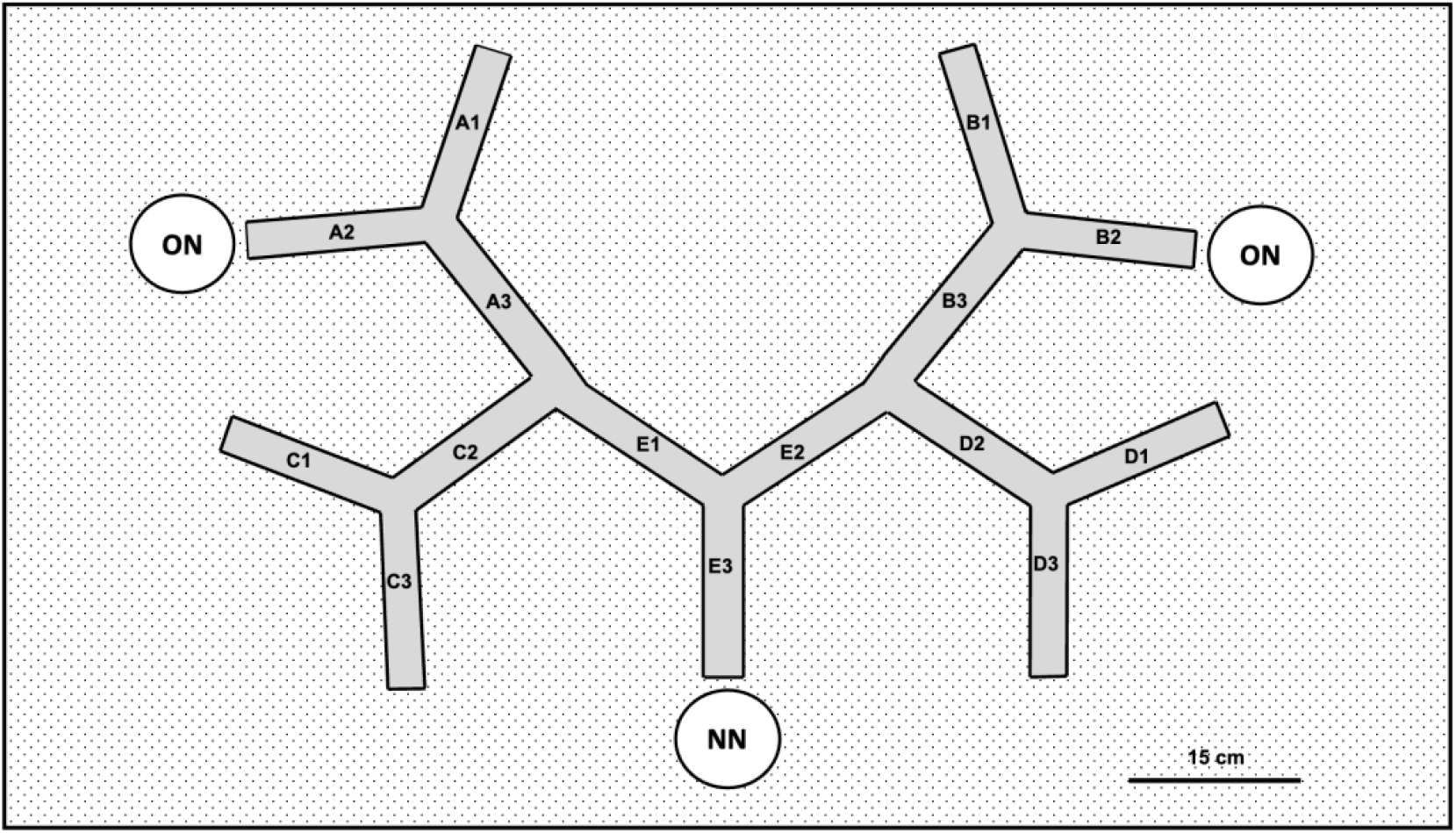
The experimental arena is 122 cm X 92 cm filled with water. The maze is placed within the arena. The maze was divided into five areas named A, B, C, D and E. Each area consists of three arms, for example area A had three arms termed as A1, A2 and A3 of length 15 cm each. Two colonies in their old nests (ON) were placed at the end of arm A2 and end of arm B2. One new nest (NN) was placed at the end of arm E3 and as seen in the figure this arm served as the common path to enter the new nest.

### Competition experiment

Two different sets of experiments were performed to understand the effect of colony size on the competition for a new resource which is a nest in our case. In the first sets of experiments, two comparable sizes of colonies (72.38 ± 29.99) and (72.76 ± 33.29) were used (Mann-Whitney test: U=144, N1=17, N2= 17, p=1). In the second sets of experiments, two colonies with non-comparable colony sizes were used termed as large (93.21 ± 14.99) and small colonies (52 ± 10.32) (Mann-Whitney test: U=45.5, N1=14, N2= 14, p < 0.05). In the course of both these experiments there was only one potential nest available and the two colonies had to compete with each other in order to occupy the new nest. After completing the experiment, the colony that successfully occupied the new nest was termed as winner colony and the colony that was unable to occupy the new nest was termed a loser colony. The total observation period for collecting both relocation and aggression interaction data was 3 hours. After this period, colonies were maintained in the arena overnight to reconfirm that the winner colony continued to occupy the new nests.

### Control 1

Ten colonies (N = 10) with 107.60 ± 47.47 adults were used in control relocation where colony members were not exposed to aggression as only a single colony is used for the relocation experiment. Thus, instead of placing two colonies for simultaneous relocation only a single colony was placed either at the end of the arm A2 or B2 randomly. The abiotic parameters like the arena size, the dimensions of the maze, water, location of old and new nest and the distance between old and new nest were identical to what has been described above (competition experiment). The initiation of relocation (by removal of roof and providing light stress) were also identical for both the sets of experiments. All other procedures regarding the collection of data are similar for both experiments.

### Control 2

Another ten colonies (N = 10) with 65.60 ± 32.42 adults were kept under regular lab conditions at 25ºC and 55% relative humidity, 12 hours of light and 12 hours dark for 7 days. These 10 colonies were not exposed to any stress factors, relocation, or aggression. All procedures regarding the collection of colonies, marking is similar across these experiments and the previous ones. These colonies were simply maintained in lab conditions in order to examine the mortality rate within the colony and compare this with the mortality observed in colonies that had to only relocate or those that had to relocate and also face aggression as they competed for new nest.

### Mortality

Mortality data were collected for all colonies that was used in competition experiment, control 1 and control 2. The number of individuals who survived and those that had died were counted and noted after 24 hours following relocation in the case of competition experiment and control 1. Even those colonies that were undisturbed in control 2 were monitored in a similar manner 24 hours after being marked and released into their new nest. The number of brood was also counted in all the colonies in each of the cases. After 7 days from the experiments (competition or relocation) mortality data was collected in a similar manner across all the replicates.

### Behavioral observation

Information regarding the relocation dynamics included-discovery time (time between the initiation of the experiment and 1^st^ individual discovering the new nest), latency (time between the first individual discovering the new nest and the first tandem pair reaching the new nest), number of discoverers (number of individuals carrying the information of new nest) and the transportation time (time between 1^st^ and last tandem pair reaching the new nest) was collected for every replicate. During relocation, sometimes the leader of one colony led the individual of another colony as a follower to the new nest, these were also noted down and termed as cross pair tandem runs. Colony relocation data like identity of tandem leaders, followers, site of initiation and site of termination and transport of any brood was decoded from the video together with the time at which it occurred. For collecting this data, a video recorder (Sony Handycam HDR-PJ230) was placed such that the events occurring on the common path (CP) and entrance of the new nests would be recorded.

Information regarding the aggression dynamics due to competition between two colonies were focused upon. The whole relocation time was divided into three parts: pre-relocation phase (time between the initiation of the experiment and the initiation of the first tandem run), during relocation phase (time between the first and last tandem run), and post-relocation phase (time between the last tandem run and the end of the three hours, i.e., the end of the experiment). Different kinds of aggressive behaviors were noted including the identities of the individuals showing aggression and that receiving the aggression together with the colony identities of each individual. (45). Two types of aggressive behaviors were observed – antennal boxing and immobilization. When two ants face each other and repeatedly touched the others’ antennae or body with their antenna we termed it as antennal boxing. This is a bi-directional interaction with both ants showing a similar action. When one ant holds the other ant and drags it or restricts the movement of that individual, we termed it as immobilization. This is a directional interaction, where in one ant was being immobilized by another and the initiator and the receiver was always clear. Aggressive interactions that occurred in the arena was recorded using All Occurrence sessions that was conducted manually. Each session lasted for 5 minutes and was followed by a 5minute gap. A total of 18 sessions that translated into 180 minutes all occurrence sessions were conducted for each experiment and the rate of aggression was collected from this information. In addition to the number of antennal boxing and immobilization (aggressive behaviours) that were recorded during these sessions, the identity of the ants involved and the location at which it occurred in the arena was also recorded for each interaction. Thus, we obtained both the temporal and spatial dynamics of aggressive behaviors. In addition to these behaviours other activities like brood theft were also noted if any. Brood theft is defined as the act of stealing the brood of one colony by a member of another colony (46). The identity of the ants involved in the stealing and the developmental stage of the brood that was stolen was noted.

### Statistical analysis

Statistical tests were performed using R (version 4.1.2). Two-tailed non-parametric tests have been used to analyze the discovery time, number of explorers, latency, colony-level aggression, number of individuals involved in showing aggression, and frequently used type of aggression by the colony members. Temporal and spatial dynamics of aggression have been analysed by the Friedmann test. Kruskal-Wallis test has been used to check the difference in mortality rates of the colonies. All the post hoc has been done by using pairwise Wilcoxon test and Dunn test. Generalized least square model has been done to check the rate of tandem run of two competing colonies over the relocation time. Generalized linear model (GLM) was used to model the outcome (Win versus Loss) using the relocation factors (discovery time, discoverers, latency and identity of colony that initiated the first tandem run) and aggression factors (time of aggression, number of aggressive events and number of individuals involved in showing aggression) as fixed factors. A zero-inflated with negative binomial distribution was used to model brood theft of comparable sizes colonies. Zero-inflated poisson model was used to model the number of brood theft by non comparable sizes of colonies. In this set of experiment, brood theft of one replicate was an outlier and therefore has been removed from subsequent analysis. The best-fitted model was chosen based on Aikaike information criterion. All the test values were regarded as significant when p < 0.05. Unless mentioned otherwise, mean ± standard deviation of all the parameters has been mentioned.

## Results

### Competition between colonies of comparable sizes

Out of 17 replicates, one colony relocated successfully into the new nest in significantly more cases (13/17), in the remaining four cases, the two competing colonies merged into the new nest. Binomial tests: Number of trials = 17; number of success: 13 and p < 0.01). The number of cross-pair tandem runs that occurred in merged cases (18.25 ± 6.58%) was higher than in the cases where there was a clear outcome in terms of win-loss (11.67 ± 8.19%). Interestingly, in all four merged cases one of the gamergates died within 7 days of the experiment. Henceforth, 13 replicates, where there was a clear outcome, have been considered for analysis.

We found that the winner colonies’ percentage of discoverers (6.02 ± 1.96%) was significantly higher as compared to the loser colony (2.81 ± 2.33%) (Mann-Whitney U test, U=151, N1=13, N2=13, p<0.01). The colony that initiated the first tandem had a significant influence on the winning the new nest (*GLM, est = 3*.*79, z = 2*.*43, p = < 0*.*05)* (For details see S2 text, table 1). Also, the rate of relocation was lower for the loser colony as compared to winner colony *(GLS, t = 4*.*71, estimate = 0*.*88, p = < 0*.*05)* (Figure 2) (For details see S2 text, table 2). The total number of aggressive behaviors shown by the winner (91.38 ± 61.72) and loser (81.84 ± 36.45) colonies were comparable (Mann-Whitney U test, U=85, N1=13, N2=13, p=1). We found that percentage of individuals involved in aggression from winner (44.77 ± 12.04%) and loser (48.32 ± 16.88%) colonies significantly impacted the win and loss of a colony (*GLM, est = - 0*.*018, z = -2*.*5, p = < 0*.*05*) (For details see S2 text, table 3). Information regarding spatial and temporal dynamics of aggression was provided in supplementary files (S1 text, S1 fig).

**Figure 2.**
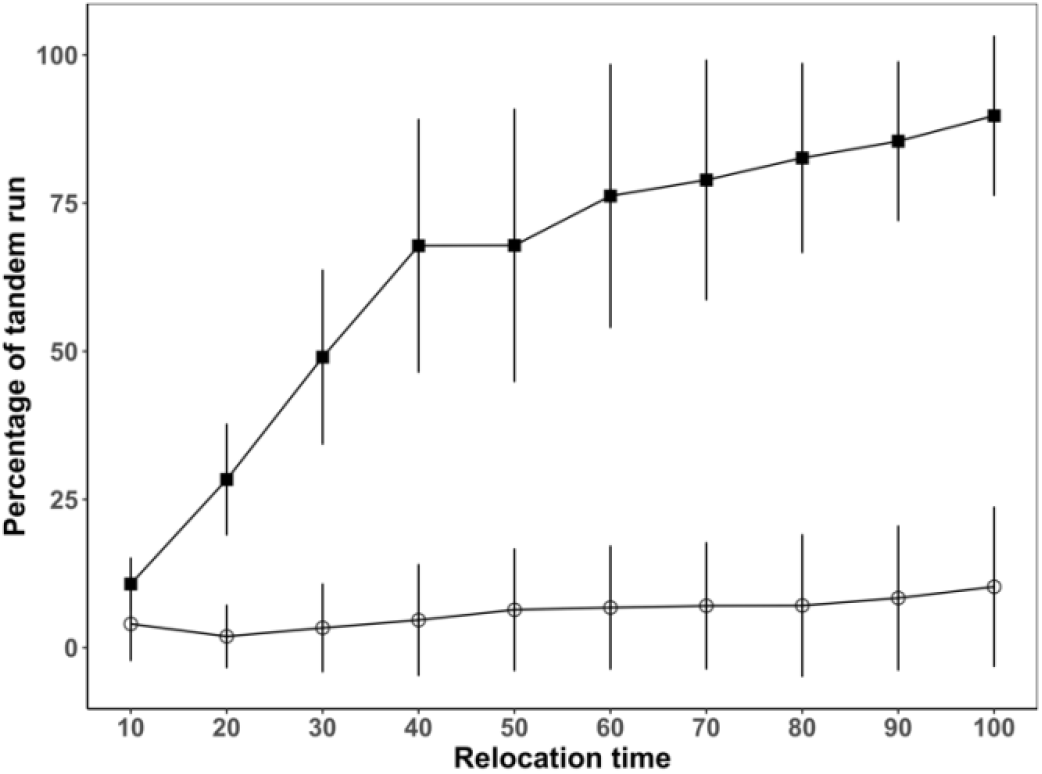
Tandem run dynamics of the winner (filled squares) and loser (open circles) colonies: The manner in which tandem runs were conducted across relocation time by the winner and loser has been represented using a line graph. The x-axis represents the percentage of relocation time divided into bins of 10% and the axis starts from the first tandem run (0%) to the last transport (100%). The y-axis represents the cumulative tandem runs by the leaders of the winning and losing colonies. The filled squares and open circles represent the average number of tandem runs over the relocation time by the winner and loser colonies respectively. The whiskers represent the standard deviation. The categories were compared using GLS, and p < 0.01 was considered statistically significant.

**Figure 3.**
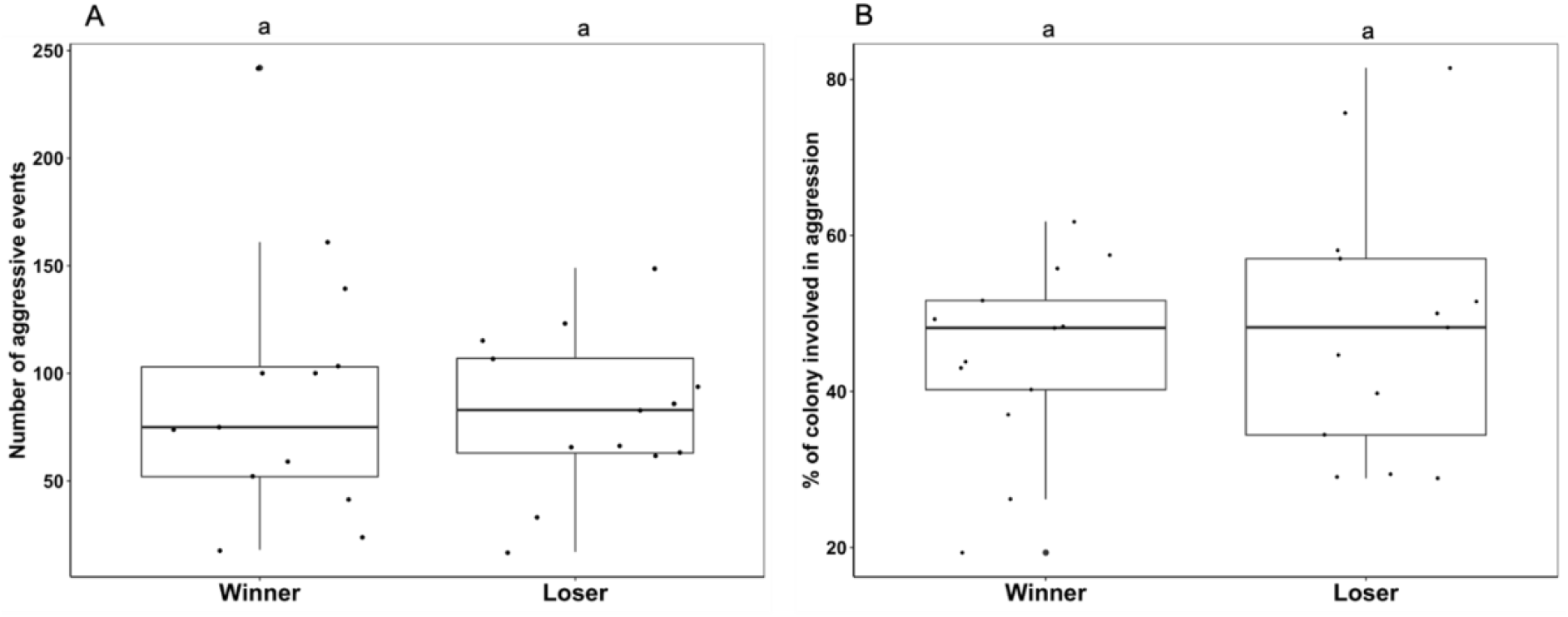
The box and jitter plot represents the aggression dynamics of the colonies. A) Total number of aggressive events shown by winner (n=13) and loser (n=13) colony. B) Percentage of winner (n=13) and loser (n=13) colony involved in aggression. The black bold line denotes the median value, the box represents the interquartile range, and whiskers represent the 1.5 times the interquartile range. The jitters denote the data points. The comparison between the two categories was made using the Mann-Whitney test. Significant differences between groups are represented by different alphabets placed above the boxes.

### Brood theft

Out of 13 replicates, brood theft occurred in 10 replicates. In 10 replicates, total of 77 brood theft events has been observed across both winner and loser colonies. As pupae was the favored item, the analysis includes only pupae theft. The number of brood theft conducted by winner colonies (3.9 ± 3.03) and loser colonies (3.8 ± 4.93) were comparable (*GLM, est = 0*.*42, z = 0*.*99, p = 0*.*3*).

### Mortality

Mortality was significantly higher for the loser colony (10.75 ± 5.39%) as compared to both control 1 and (3.04 ± 1.85%) and control 2 (1.91 ± 2.22%) and the mortality of the winner colony (8.99± 8.83%) was significantly higher as compared to control 1 after 24 hours of observation (Kruskal-Wallis chi-squared test: 21.40, df=3 p < 0.05) (Figure 4A). After 7 days, the mortality was significantly higher for the loser colony (23.44 ± 12.40%) than control 2 (8.29 ± 6.33%). The mortality of the winner colony (18.57 ± 10.35%) was comparable with control 1 (13.16 ± 8.04%), control 2 and loser colony (Kruskal-Wallis chi-squared test: 12.18, df = 3 p < 0.05) (Figure 4B). All the post hoc results have been provided in supplementary files (For details see table S2 text, 4A, 4B).

**Figure 4.**
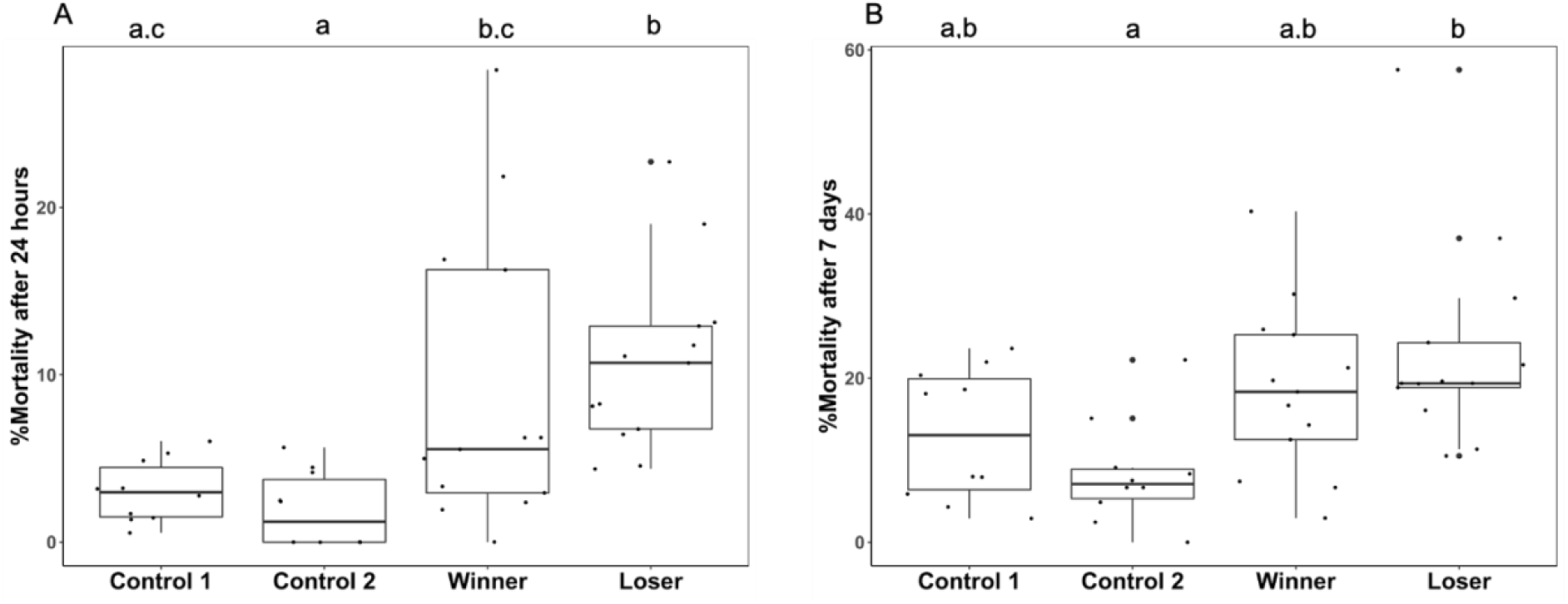
The box and jitter plots represent %mortality across experiments. A) %Mortality of control 1 (colonies expose to relocation, control 2 (laboratory condition) winner and loser colonies after 24 hours B) %Mortality of control 1 (colonies expose to relocation), control 2 (laboratory condition) winner and loser colonies after 7 days. The black bold line denotes the median value, the box represents the interquartile range, and whiskers represent the 1.5 times the interquartile range. The jitters denote all the data points. The comparison between the categories was made using the Kruskal-Wallis chi-squared test. Significant differences between groups are represented by different alphabets placed above the boxes.

### Competition between colonies with non-comparable colony sizes

Out of 14 replicates, one of the competing colonies relocated successfully in 11 cases. In the remaining three cases, the two competing colonies merged into the new nest (Binomial tests: Number of trials = 14; Number of success = 11 and p < 0.05). In these three cases, colonies merged post relocation. Number of cross pair tandem runs that occurred in merged cases (24.85 ± 18.51%) was higher than in the cases where only one colony relocated to the new nest (2.51 ± 3.18%). In all three merged cases, one of the gamergate died within 7 days of the experiment. Out of 11 cases, the larger colony relocated successfully in 10 cases (Binomial tests: Number of trials = 11; Number of success = 10 and p < 0.05). The proportion of replicates where larger colonies relocated successfully termed as winner colony is significantly higher than the proportion of colonies where smaller colonies termed as loser colony relocated and both colonies merged. Hence, colony size has a major impact on the occupancy of the new nest. Further analysis has been performed with those 10 replicates, where the larger colony was the winner. We found that the percentage of discoverers of winner colonies (8.0 ± 4.55%) was significantly higher as compared to the loser colony (2.43 ± 2.80%) (Mann-Whitney U test, U=88, N1=10, N2=10, p<0.05). Moreover, the colony that initiated the first tandem running had a major influence on the occupancy by that particular colony in the new nest (*GLM, est = 3*.*59, z = 2*.*30, p = < 0*.*05)* (For details see S2 text, table 5). Aggression per unit time at the location E (near to the new nest) *(GLM: est = 48*.*06, z = 2*.*9, p < 0*.*05)* and the percentage of individuals of winner (40.79 ± 9.93) and loser (63.71 ± 16.85) colony showing aggression *(GLM: est = -0*.*16, z = -7*.*53, p < 0*.*05)* had an impact on the outcome (For details see S2 text, table 6). The total number of aggressive behavior shown by the winner (116.8 ± 39.13) and loser (111.3 ± 31.38) colonies were comparable (Mann-Whitney U test, U=85, N1=13, N2=13, p=1). Information regarding spatial and temporal dynamics of aggression was provided in supplementary files (S1 text, S2 Fig).

**Figure 5.**
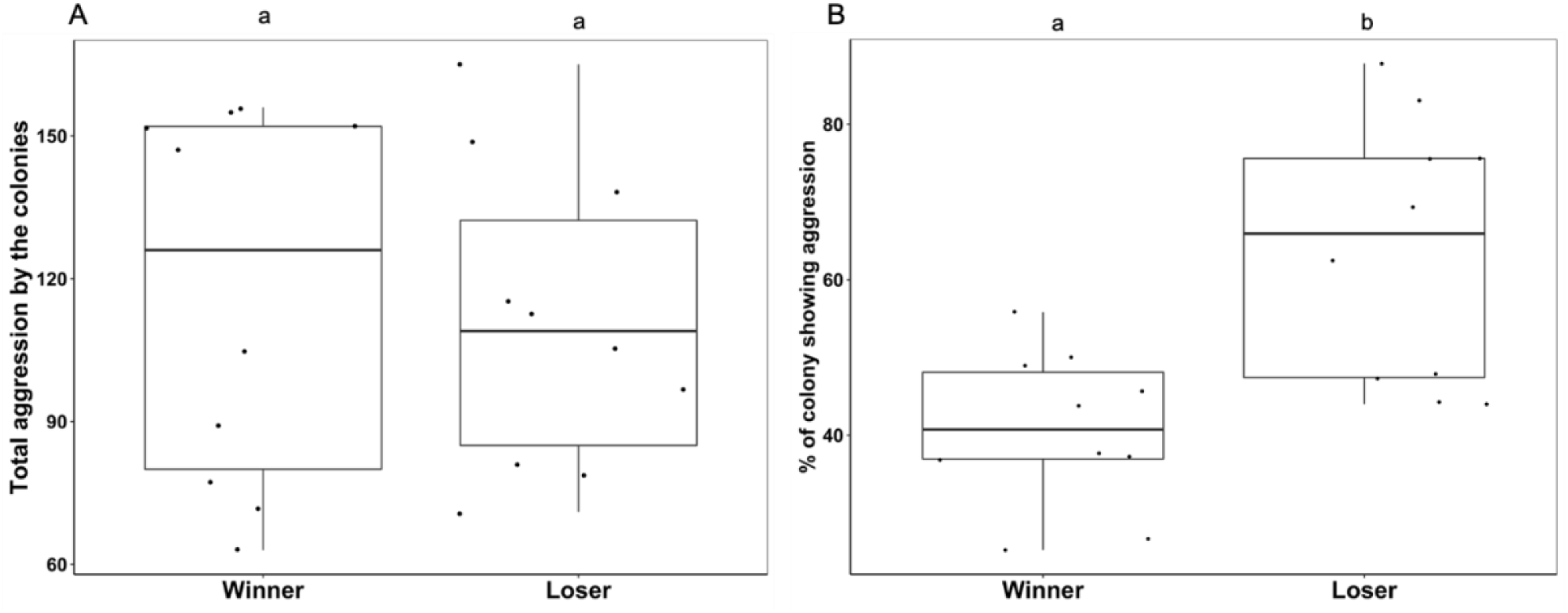
The box and jitter plot represents total number of aggressive events shown by winner (W) and loser (L) colony. The black bold line denotes the median value, the box represents the interquartile range, and whiskers represent the 1.5 times the interquartile range. The jitters denote all the data points. The comparison between the two categories was made using the Mann-Whitney test. Significant differences between groups are represented by different alphabets placed above the boxes.

### Brood theft

Out of 10 replicates, brood theft occurred in 9 replicates. In 9 replicates, total of 45 brood theft events has been observed including both winner and loser colonies. The number of brood theft (pupae theft) done by winner colonies (2 ± 2.3) was significantly higher than loser colonies (0.25± 0.4) (*GLM, est = -2*.*75, z = -3*.*65, p < 0*.*05*).

### Mortality

We have found that the mortality was significantly higher for the loser colony (8.05 ± 4.19%) as compared to both the control 1 (3.04 ± 1.85%) and control 2 (1.91 ± 2.22%) and the mortality of the winner colony (8.35 ± 5.50%) was significantly higher as compared to control 2 after 24 hours of observation (Kruskal-Wallis chi-squared test: 17.89, df = 3 p < 0.05) (Figure 6A). After 7 days, mortality was significantly higher for the winner (26.76 ± 20.05%) and loser (26.42 ± 15.16%) colony than control 2 (8.29 ± 6.33%) and comparable with control 1 (13.16 ± 8.04%) (Kruskal-Wallis chi-squared test: 13.75, df = 3 p < 0.05) (Figure 6B). All the post hoc results were mentioned on supplementary files (For details see table S2 text, 7A, 7B).

**Figure 6.**
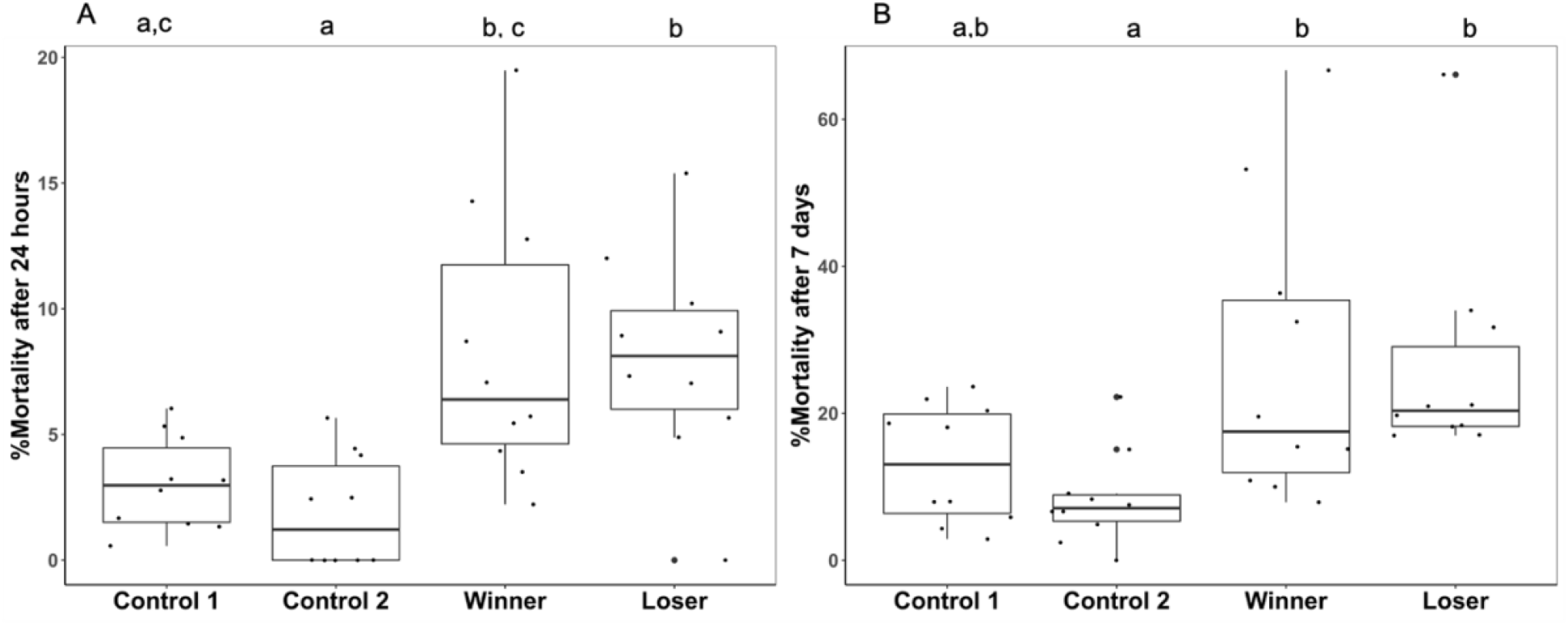
The box and jitter plot represents %mortality of control 1, control 2, large and small colonies. A) Percentage mortality of control 1 (colonies expose to relocation), control 2 (laboratory condition) large and small colonies after 24 hours b) Percentage mortality of control 1 (colonies expose to relocation), control 2 (laboratory condition) large and small colonies after 7 days. The black bold line denotes the median value, the box represents the interquartile range, and whiskers represent the 1.5 times the interquartile range. The jitters denote all the data points. The comparison between the categories was made using the Kruskal-Wallis chi-squared test in R. Significant differences between groups are represented by different alphabets placed above the boxes.

## Discussion

Competition not only directly or indirectly affects how species gain access to the resources it also impacts the species composition of a region. Resource allocation, colony fission, and territoriality are major causes of intraspecific competition in ants. For example, the degree of intraspecific competition in *Aphaenogaster serilis* and *Cataglyphis cursor* impacts access of resources and reduction of their colony size. (47,48). Some species like *Camponatus cruentatus* and *Messor aciculatus* show ritualized combat and intracolonial aggression respectively over the conspecific colonies (10,49). Interestingly, due to intraspecific competition, *Pheidole pallidula* increase their soldier production and *Formica altipetens* increase their nest size and the distance from their nearest neighbours (50,51). In the current study, we have examined intraspecific competition in the context of relocation which is a goal-oriented task involving the whole colony with the new nest as a limited resource, for the first time to the best of our knowledge. Such intraspecific competition is expected to occur when a large area hosting multiple colonies gets disturbed due to large-scale events like monsoons in the natural habitat. Thus, this lab study reflects a phenomenon that is likely to occur in the natural habitat.

Here, we used the nest as a limiting resource and two colonies of *Diacamma indicum* were motivated to relocate from their old nest to a new nest. In 13/17 trials, one colony successfully relocated to a new nest when colonies of comparable sizes were competing with each other. When non comparable sizes of colonies were competing, in 11/14 trials, we had a clear outcome. Interestingly, in approximately 25% of the cases, both colonies merged together and the gamergate of one of the colonies was found dead leading to a single reproductive individual within the colony. When non comparable colonies were competing, the gamergate of the smaller colony died. Further investigation into factors like the age of the gamergate, its fecundity and aggression need to be conducted to understand the reason for the demise of one of the gamergate in merging colonies. Colony merger was brought about by the significant increase in cross pair tandem running, cases where a tandem leader from one colony was followed by ants from another colony. On average 1.56 times increase in cross pair tandem running for comparable size colonies and 9.9 times increase in non-comparable size colonies was observed in the merged cases. This is another feature that needs further exploration. Do tandem leader relax their threshold for nestmate specific behaviours when they are actively involved in transportation, is a specific question that needs to be explored. This merger of colonies has major implications for the relatedness between the colony members and has been discussed together with one other phenomenon, brood theft that is also likely to impact relatedness. Ants have the tendency to steal brood from foreign colonies as it plays an important role in the amelioration of their colonies. It acts as a food source, especially larvae (16). It is reported in several contexts like obtaining nutrition, increasing the workforce and the survivability of the colonies (16,52). Brood theft has been previously studied in the temperate regions in several ant genera such as *Solenopsis, Myrmecocystus, Polyergus, Strongylognathus* (53). *Solenopsis invicta* and *Myrmecosystus mimicus* are well known for intraspecific brood raiding from the nearby colonies which leads to an increases in the mortality of incipient nests (54,55). Brood theft was reported in *Diacamma indicum* tropical ponerine ant (56). Pupae are the most preferred item, this does not need a large investment in terms of food and care to eclose and increase the workforce as it’s the last developmental stage (57). In the current study, stealing brood increased when colonies were exposed and were in the process of relocation. When two similar colony sizes were relocating and competing with each other for the new nest, both colonies did comparable number of stealing. However, when two dissimilar sizes of colonies were competing, the mean number of successful stealing by larger colonies was 8 times higher than smaller colonies. On average 39.05 ± 39.97% brood was stolen indicating that this is a relatively large perturbation to the colony. In the natural habitat, similar levels of pupa theft were recorded. Brood theft and colony merger together would reduce the relatedness between colony members from the theoretical expected value of 0.75 for colonies having a single reproductive that has mated with only a single male. Given these observations, it becomes essential to check the actual relatedness values of this species in future studies and pounder about the reasons for cooperation in this species.

In order to delineate the factors that determine which colony wins the resource – new nest in this case we conducted several analyses. We found that when two colonies of comparable sizes were competing with each other, colonies that had more members discovering the new nest and colonies that initiated the first tandem run has a greater chance of occupying the new nest and outcompeting another colony. Subsequently, the winning colonies showed a higher rate of relocation into the new nest. If the colonies of unequal sizes were competing, then the smaller colony was unable to attain the resource. Discovery of resource is an important component of the competition dynamics regarding food in other ant species. In the case of woodland ants like *Camponatus ferrugineus, Lasius alienus, Formica subsericea* and *Prenolepis imparis* studies demonstrate a trade-off between resource discovery and behavioral dominance of resources (12, 51). If the group size is not similar and they compete for the same resource, then the smaller colony was at a disadvantage as members engaged in interactions with the competitors could not simultaneously undertake the assigned tasks (59). In the current study, we hypothesize that smaller colonies were not able to compete for the resource because their workforce was mostly occupied with interactions with larger colonies and could not allocate members for exploration leading to delayed discovery, higher latency to initiate tandem run. During intraspecific competition alien conspecific are known to receive aggression and the degree of aggression is typically context-dependent (60). Previous studies not only show that larger colonies display more aggressive interaction but also that there is a positive correlation between individual levels of aggression and colony size in *Leptothorax ambiguous* in the context of nestmate recognition and territoriality (61). However, there was no such relationship between individual aggression and colony size in *Rhytidoponera confuse* when they encountered conspecifics in the context of nestmate recognition (62). In current study, we found that irrespective of colony size and resource dominance, the aggression level across both the colonies involved was comparable. Aggressive interactions did not significantly influence colonies chances of winning resources. Although, the rate of aggression was much higher and more intense towards the vicinity of their old nest and the new nest. probably because they wanted to guard their brood and nest (63).

Many worker ants were killed due to competition. Mortality due to competition was a threat to the ant colonies. Competition can be affected by many factors like natural and anthropogenic disturbances and temperature (64). In our study, ant colonies underwent two kinds of stress simultaneously: relocation and competition as two colonies compete and fight with each other for a nest during relocation. We found that the process of relocation imposed significantly higher mortality as compared to controls that were not involved in relocation. Relocating colonies on average showed 3.68 fold increase across 24 hours and 2.44 fold increase across 7 days as compared to colonies that did not participate in relocation. Competition for the resource and ensuing aggression significantly increased the mortality among the losing colonies over the short term that is across 24 hours, even though the magnitude of this was not very high.

In conclusion, examining competition for new nest is essential for understanding the structure and composition of ant communities. Our study indicates that it is also essential to examine competition for new nest in order to understand the basic features of the colony like the relatedness between nestmates and mortality among these superorganisms.

## Acknowledgment

We thank Mr. Basudev Ghosh for his assistance and maintenance of *Diacamma indicum* colonies. We thank Dr. Rubina Mondal for her valuable input in statistical analysis. We also thank Indian Institute of Science Education and Research Kolkata for the support they have given and Science and Engineering Research Board (SERB), Govt. of India for EMR/2017/00147.

