## Supplementary files for "Intraspecific competition for a nest and its implication for the fitness of relocating ant colonies"

**Supplementary Material**

^1^Department of Biological Sciences, Indian Institute of Science Education and Research Kolkata, Mohanpur, West Bengal, 741246, India

**^1*^Address for correspondence:**

Behaviour and Ecology Lab, Department of Biological Sciences, Indian Institute of Science Education and Research Kolkata, Mohanpur, West Bengal 741246, India

**Figure S1: Spatial and temporal dynamics of aggression**

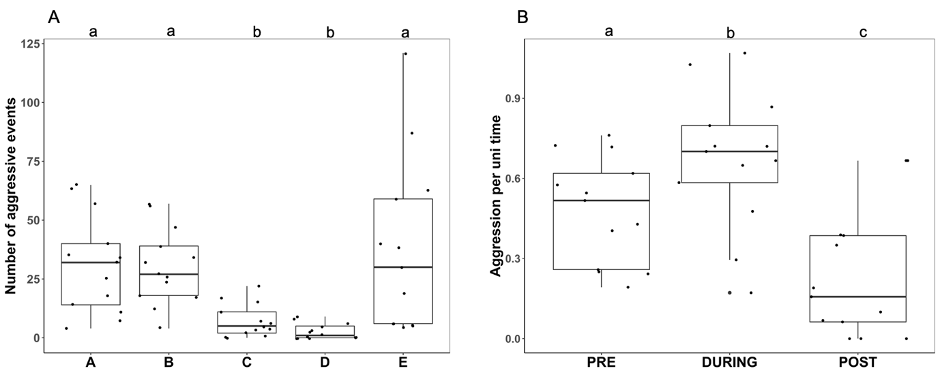

**Aggression dynamics for comparable size colonies**

We have found that total aggressive interaction was significantly higher towards the two old nests and one new nest compared to other parts of the arena (Friedmann test X^2^: 36.92, df = 4, p < 0.05) (S1A fig). We found that total aggressive interaction over the relocation time was significantly different across three phases (Friedmann test X^2^: 20.462, df = 2, p < 0.05) (Figure S1B Fig). The post hoc results have been provided in the supplementary table (Table 1A and 1B). Hence, more aggressive events occurred toward the old and new nest during the relocation phase.

**Figure S2: Aggression dynamics for non-comparable size colonies**

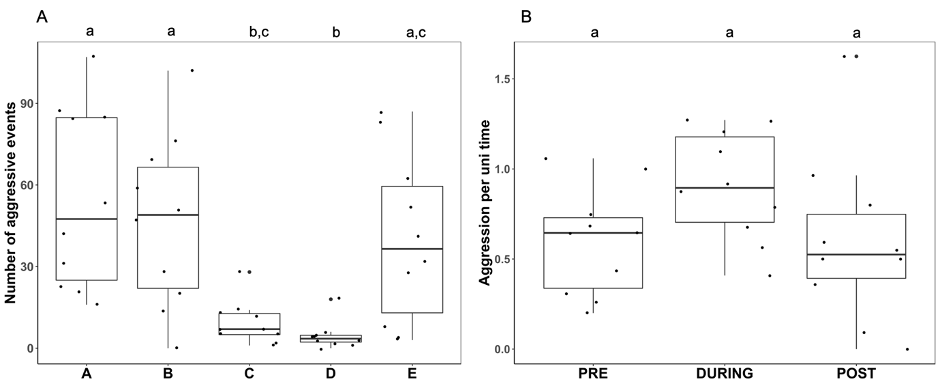

In this set of experiments, we found that total aggressive interactions were significantly higher towards the old and new nests (Friedmann test X^2^: 23.81, df = 4, p < 0.05) (S2A Fig). By analyzing the aggression dynamics of the three phases of relocation, we have found that overall aggression dynamics across the three phases was comparable (Friedmann test X^2^: 5.6, df = 2, p = 0.06) (S2B Fig). The post hoc results have been provided in the supplementary table (S2 text, Table 2).

**Table 1A. Details of post-hoc results of total number of aggressive events that occurred in all five locations across the maze.**

|  | **A** | **B** | **C** | **D** |
| --- | --- | --- | --- | --- |
| **B** | 1.000 |  |  |  |
| **C** | <0.05 | <0.05 |  |  |
| **D** | <0.05 | <0.05 | 0.082 |  |
| **E** | 1.000 | 1.000 | <0.05 | <0.05 |

**Table 1B . Details of post-hoc results of total number of aggressive events over the relocation time.**

|  | **Pre** | **During** |
| --- | --- | --- |
| **During** | <0.05 |  |
| **Post** | < 0.05 | <0.05 |

**Table 2. Details of post-hoc results of total number of aggressive events that occurred in all five locations across the maze.**

|  | **A** | **B** | **C** | **D** |
| --- | --- | --- | --- | --- |
| **B** | 1.000 |  |  |  |
| **C** | <0.05 | <0.05 |  |  |
| **D** | <0.05 | <0.05 | 0.29 |  |
| **E** | 0.97 | 1.000 | 0.14 | <0.05 |

**S2 Table: GLM and GLS Results of Relocation dynamics and Post hoc results of Mortality of Winner and Loser colonies (Comparable and non comparable colonies)**

**Table 1: Details of GLM analysis to investigate the relocation parameters impacting the win/loss of a colony while competing for the new nest.**

Model <- glm (Win/loss ~ Initiation of tandem run + Latency, data= Relocation_GLM, family= binomial)

|  | **Estimate** | **Std. Error** | **z value** | **p value** |
| --- | --- | --- | --- | --- |
| **Intercept** | -0.54 | 1.77 | -0.30 | 0.75 |
| **Initiation of tandem run** | 3.79 | 1.55 | 2.43 | < 0.05 |
| **Latency** | -0.02 | 0.03 | -0.74 | 0.45 |

The response “Win/loss” is the final outcome of the colony in terms of the occupancy of new nest. The fixed effect is “Initiation of tandem run” which is the colony that initiated the first tandem run during relocation. The fixed effect “Latency” is the time between the discovery of a new nest and the initiation of the tandem run of each colony.

**Table 2: Details of GLS analysis to compare the progression of number of tandem run over the relocation time in winner and loser colony**.

Model <- gls (Percentage of tandem run ~ Winner/Loser * Relocation time, correlation = corAR1( form = ~ 1|ColonyID), data= TR_Time_GLS, method = “ML”)

| **Coefficients** | | | | |
| --- | --- | --- | --- | --- |
|  | **Value** | **Std. Error** | **t value** | **p value** |
| **Intercept** | -0.41 | 0.26 | -1.5 | 0.12 |
| **Relocation time** | 0.10 | 0.03 | 2.94 | <0.05 |
| **Winner** | -4.41 | 1.18 | -3.73 | <0.05 |
| **Winner:Time** | 0.88 | 0.18 | 4.71 | <0.05 |

The response variable “% tandem run” is the percentage of tandem run over the relocation time, the fixed effect “Winner/Loser” is the type of colony that wins and lose, and the fixed effect “Relocation time” is the percent of time elapsed in bins of 10%. The identity of the colonies used is used as the grouping factor “ColonyID”.

**Table 3: Details of GLM analysis to investigate the aggression dynamics impacting the win/loss of a colony while competing for the new nest.**

Model <- glm (Win/loss ~ Interaction + Percentage of individual; data= Agg_GLM, family= binomial)

|  | **Estimate** | **Std. Error** | **z value** | **p value** |
| --- | --- | --- | --- | --- |
| **Intercept** | 0.83 | 0.35 | 2.34 | < 0.05 |
| **Interaction** | 2.04 | 2.58 | 0.79 | 0.42 |
| **Percentage of individual** | -0.018 | 0.007 | -2.52 | < 0.05 |

The response “Win/loss” is the final outcome of the colony in terms of the occupancy of new nest. The fixed effect is “Interaction” which is the number of aggression interactions at a particular location. The fixed effect “Percentage of individual” is the percentage of individuals involved in aggression.

**Table 4A. Details of post-hoc results of mortality after 24 hours between controls , winner and loser colonies.**

| **Category** | **P value** |
| --- | --- |
| **Control 2 – Control 1** | 0.41 |
| **Control 2 – Loser** | < 0.05 |
| **Control 1 – Loser** | < 0.05 |
| **Control 2 – Winner** | < 0.05 |
| **Control 1 – Winner** | 0.19 |
| **Losing - Winner** | 0.22 |

“Control 1” are the colonies that are exposed only to relocation. “Control 2” are the colonies that are not exposed to either relocation or competition. Wining colonies are colonies that are successful to occupy the new nest. Losing colonies are the colonies that are unable to occupy the new nest.

**Table 4B. Details of post-hoc results of mortality after 7 days between controls, winner and loser colonies.**

| **Category** | **P value** |
| --- | --- |
| **Control 2 – Control_1** | 0.74 |
| **Control 2 – Loser** | < 0.05 |
| **Control_1 – Loser** | 0.16 |
| **Control 2 – Winner** | 0.07 |
| **Control_1 – Winner** | 0.7 |
| **Losing - Winner** | 0.36 |

“Control 1” are the colonies that are exposed only to relocation. “Control 2” is the colonies that are not exposed to either relocation or competition. Wining colonies are colonies that are successful to occupy the new nest. Losing colonies are the colonies that are unable to occupy the new nest.

**Table 5: Details of GLM analysis to investigate the relocation parameters impacting the larger colony to win while competing for the new nest.**

Model <- glm (Win/loss ~ Percentage of discoverers + Initiation of tandem run, data= GLM_Relocation, family= binomial)

|  | **Estimate** | **Std. Error** | **z value** | **p value** |
| --- | --- | --- | --- | --- |
| **Intercept** | -3.10 | 1.52 | -2.04 | < 0.05 |
| **Percentage of discoverers** | 0.27 | 0.26 | 1.06 | 0.28 |
| **Initiation of tandem run** | 3.59 | 1.55 | 2.30 | < 0.05 |

The response “Win/loss” is the final outcome of the colony in terms of the occupancy of new nest. The fixed effect is “Initiation of tandem run” which is the colony that initiated the first tandem run during relocation. The fixed effect “Percentage of discoverers” is the number of individuals who -discovered the new nest.

**Table 6: Details of GLM analysis to investigate the aggression dynamics impacting the wining of a larger colony while competing for the new nest.**

Model <- glm (Win/loss ~ Location * Aggression/Time + Percentage of individual; data= Agg_GLM, family= binomial)

|  | **Estimate** | **Std. Error** | **z value** | **p value** |
| --- | --- | --- | --- | --- |
| **Intercept** | 8.14 | 1.14 | 7.14 | < 0.05 |
| **Location B** | -0.14 | 0.65 | -0.22 | 0.82 |
| **Location C** | -0.06 | 0.63 | -0.10 | 0.91 |
| **Location D** | -0.22 | 0.63 | -0.35 | 0.72 |
| **Location E** | -1.15 | 0.67 | -1.71 | 0.08 |
| **Aggression/Time** | 1.52 | 5.50 | 0.27 | 0.78 |
| **Percentage of individual** | -0.16 | 0.02 | -7.53 | < 0.05 |
| **Location B : Aggression/Time** | 2.31 | 6.71 | 0.34 | 0.73 |
| **Location C : Aggression/Time** | 32.65 | 46.19 | 0.70 | 0.47 |
| **Location D : Aggression/Time** | 56.98 | 33.12 | 1.72 | 0.08 |
| **Location E : Aggression/Time** | 48.06 | 16.56 | 2.90 | < 0.05 |

The response “Win/loss” is the final outcome of the colony in terms of the occupancy of new nest. The fixed effect is “Location” which is the location where aggression is taking place. The fixed effect is “Aggression/Time” is the number of aggression taking place at aprticular location at per unit time. The fixed effect is “Percentage of individual” is the percentage of individual involved in aggression.

**Table 7A. Details of post-hoc results of mortality after 24 hours between controls , larger and smaller colonies.**

| **Category** | **P value** |
| --- | --- |
| **Control 2 – Control 1** | 0.7 |
| **Control 2– Winner** | < 0.05 |
| **Control 1 – Winner** | 0.053 |
| **Control 2 – Loser** | < 0.05 |
| **Control 1 – Loser** | < 0.05 |
| **Winner - Loser** | 0.8 |

“Control 1” are the colonies that are exposed only to relocation. “Control 2” is the colonies that are not exposed to either relocation or competition. Larger colonies are colonies that are successful to occupy the new nest. Smaller colonies are the colonies that are unable to occupy the new nest.

**Table 7B. Details of post-hoc results of mortality after 7 days between controls, larger and smaller colonies.**

| **Category** | **P value** |
| --- | --- |
| **Control 2 – Control 1** | 0.3 |
| **Control 2– Winner** | < 0.05 |
| **Control 1 – Winner** | 0.39 |
| **Control 2 – Loser** | < 0.05 |
| **Control 1 – Loser** | 0.15 |
| **Winner - Loser** | 0.57 |

“Control_1” are the colonies that are exposed only to relocation. “Control 2” is the colonies that are not exposed to either relocation or competition. Larger colonies are colonies that are successful to occupy the new nest. Smaller colonies are the colonies that are unable to occupy the new nest.
